# Evaluation of *Leishmania* Homologue of Activated C Kinase (LACK) of *Leishmania donovani* in comparison to glycoprotein 63 as vaccine candidate against visceral leishmaniasis

**DOI:** 10.1101/2025.06.25.661467

**Authors:** Nicky Didwania, Sudipta Bhowmick, Abdus Sabur, Anirban Bhattacharya, Nahid Ali

**Author notes:** **To whom correspondence should be addressed:** Prof. Nahid Ali / /5197.

## Abstract

Leishmaniasis, caused by *Leishmania* protozoa transmitted via sand fly bites, affects over 12 million people annually, manifesting as self-limiting cutaneous lesions or fatal visceral leishmaniasis (VL). The disease’s immune response involves a Th1/Th2 paradigm, with Th1 promoting resistance and Th2 linked to susceptibility. Despite no available human vaccine, the *Leishmania* homologue of activated C kinase (LACK) protein has shown promise as a candidate due to its conservation across species. In this study, we purified native 34-kDa LACK protein from *Leishmania donovani* promastigotes and compared its efficacy with gp63 in cationic DSPC liposomes. While gp63 exhibited protective efficacy, LACK failed to protect BALB/c mice. Recombinant LACK, which was cloned, expressed, and purified, also did not confer protection. Immunological assays revealed a Th2-biased immune response characterized by a high IgG1/IgG2a ratio, elevated Th2 cytokines, and an unaltered delayed-type hypersensitivity (DTH) response, highlighting its limited potential.

**Importance:** This study underscores the limitations of LACK as a standalone vaccine candidate against VL. Both native and recombinant LACK forms, despite liposomal encapsulation, elicited a Th2- dominated immune response, inadequate for protection. These findings suggest that LACK’s intrinsic immunogenic properties may not support the robust Th1 response crucial for combating *Leishmania donovani* infections. Future research should prioritize alternative antigens or combination formulations to promote Th1-driven immunity. This work contributes to the growing body of knowledge guiding vaccine development for VL, emphasizing the need for immunogens that effectively target protective immune pathways.

## INTRODUCTION

Leishmaniasis, caused by protozoan parasites of the *Leishmania* genus, presents in various forms, with visceral leishmaniasis (VL) being the most severe. Known as kala-azar or “black fever,” VL is a major global health threat, endemic in over 70 countries and putting more than 1 billion people at risk. An estimated 50,000 to 90,000 new VL cases occur annually, with the majority reported from Brazil, Ethiopia, India, Kenya, Somalia, South Sudan, and Sudan (1). The emergence of post-kala-azar dermal leishmaniasis (PKDL) has further exacerbated concerns. Though PKDL has a lower mortality rate than VL, it has significant socioeconomic consequences, as patients serve as reservoirs for parasites, perpetuating transmission and potentially causing new VL cases (2). VL treatment predominantly relies on chemotherapy, but rising drug resistance, toxicity, and high hospitalization costs pose major challenges. These limitations underscore the urgent need for safer, more effective alternatives. Vaccination offers a promising prevention strategy, potentially providing immunity superior to that of treatment. Despite considerable research into vaccine development, no vaccine for human leishmaniasis has been approved to date. In recent years, significant progress has also been made in clinical-stage *Leishmania* vaccine development. For instance, the adenoviral vectored vaccine ChAd63-KH14, encoding two *Leishmania* antigens and developed by Paul Kaye’s group, has shown safety and immunogenicity in early-phase clinical trials (3, 4). Another major development is the live attenuated *Leishmania major* centrin gene-deleted vaccine (*L. major* cen−/−), pioneered by Hira Nakhasi’s group, which has demonstrated promising protective efficacy in preclinical models (5). These candidates represent important milestones in the translational pipeline and provide a framework for evaluating new vaccine formulations and antigens.

Researchers have explored several vaccine strategies based on insights from leishmanization and its protective immune responses, including the use of whole-killed or attenuated parasites, recombinant proteins, and DNA vaccines (6). One of the most promising candidates is the *Leishmania* homologue of activated C kinase (LACK), a 34-kDa protein conserved across *Leishmania* species and life stages. LACK is crucial for various cellular processes essential to leishmanial infection (7). Initial studies indicated that LACK could serve as a potential vaccine target (8). Mice expressing LACK in their thymus exhibited reduced Th2 responses and a healing phenotype, suggesting its potential for inducing immune tolerance. Furthermore, LACK- specific T-cell expansion supported its candidacy as a vaccine target. DNA vaccines encoding the LACK gene have demonstrated protective effects in mice against *L. major*, particularly when combined with viral vectors as boosters (9) or fused with other antigens such as TSA (10). Intranasal administration of LACK DNA has induced protective responses in mice and hamsters, offering protection against both cutaneous and visceral leishmaniasis (11–13).

However, immunization with the LACK protein alone failed to protect BALB/c mice against *Leishmania amazonensis* (14). Similarly, LACK as a peptide antigen showed limited efficacy in cutaneous leishmaniasis (CL), despite inducing high IFN-γ levels (15). Nonetheless, recent studies continue to explore LACK in updated vaccine platforms. For example, a non-replicative DNA vaccine encoding LACK demonstrated significant efficacy against canine VL caused by *L. infantum*, suggesting that under optimized conditions, LACK may retain immunological potential (16). Combining LACK with IL-12, yielded promising results in protecting against *Leishmania* species. Oral administration of genetically engineered *Lactococcus lactis* expressing both recombinant LACK and IL-12 successfully induced mucosal immunity and protected mice against *L. major* challenges (17). Additionally, a multiepitope vaccine comprising LACK, LeIF, GP63, and SMT antigens from *L. major* demonstrated partial immunity (18). A heterologous prime-boost approach, using a non-replicative vaccinia recombinant vector expressing LACK, provided protection against both canine and murine VL. This strategy induced a predominantly Th1-specific immune response, showing potential as a viable vaccine candidate (19, 20).

The development of an effective vaccine for VL requires a comprehensive understanding of VL immunology and the factors influencing protective immune responses (21). Evidence from live vaccines and naturally acquired immunity suggests that long-lasting protection against reinfection is possible. Persistent antigens or pathogens are essential for the expansion of multifunctional CD4+ and CD8+ T cells, which enhance both effector and memory responses (22, 23). Therefore, a balanced approach is necessary to minimize the toxicity associated with live vaccines while optimizing the immunogenicity and efficacy of defined or subunit vaccines (24). While LACK has been previously investigated using DNA-based and recombinant protein vaccine strategies, these earlier studies employed conventional adjuvants such as alum or incomplete Freund’s adjuvant, which have limited ability to promote cross-priming and durable cell-mediated responses. In contrast, DSPC liposomes have demonstrated potent adjuvanticity and the capacity to generate robust CD4+ via MHC Class II and facilitate antigen entry into the MHC Class I pathway effectively inducing CD8+ T cell, which are essential for memory responses and long lasting immunity as depicted in our earlier work (25). Notably, to the best of our knowledge, recombinant LACK protein has not been tested in conjunction with this liposomal delivery platform in a BALB/c model of VL. The present study therefore sought to determine whether this optimized delivery system could overcome the intrinsic limitations of

LACK observed in previous studies or reaffirm its inadequacy as a standalone vaccine antigen. This approach allows a definitive evaluation of LACK’s potential within a state-of-the-art immunopotentiating system and provides valuable comparative insights relative to a benchmark protective antigen gp63 (26). Based on these observations, we aimed to evaluate the efficacy of a 34-kDa protein purified from *Leishmania donovani* promastigotes (LAg), encapsulated in DSPC liposomes and compared its efficacy to a protective antigen, gp63. This protein was identified by MALDI-TOF as the LACK antigen. To rigorously assess the immunogenicity and protective efficacy of LACK, we cloned, expressed, and purified recombinant LACK protein and entrapped it in DSPC-based cationic liposomes for immunization studies in BALB/c mice.

Thus, the present study does not aim to revive LACK uncritically, but rather to provide a definitive evaluation of its vaccine potential under optimized liposomal conditions, where other antigens (e.g., gp63) have shown strong protection. This strategy allows us to isolate antigen quality as a variable and draw more reliable conclusions about its candidacy.

## RESULTS

### Physicochemical Characterization of Liposomes

The mean particle size and surface charge of the prepared liposomes were evaluated by dynamic light scattering and zeta potential analysis, respectively. Liposomal formulations exhibited sizes within the 200–250 nm range and high positive surface charges, consistent with their cationic nature. Specifically, liposomal LACK showed a size of 215.46 ± 8.1 nm and a zeta potential of 51.8 ± 7.05 mV, while similar profiles were observed for Liposomal GP63 and Empty Liposome (Table 1).

**Table 1.** Physicochemical characterization of liposomal formulations and unloaded vesicles.

| Serial No | Name | Size (nm) | Charge (mV) |
| --- | --- | --- | --- |
| 1 | Liposome | 272.8± 8.8 | 51.5 ± 1.25 |
| 2 | gp63 in Liposome | 276.76± 6.9 | 51.0 ± 7.9 |
| 3 | LACK in Liposome | 215.46 ± 8.1 | 51.8 ± 7.05 |

### gp63 but not 34-kDa antigen was protective against *L. donovani* AG83

LAg proteins were separated using 10% SDS-PAGE, and both gp63 and the 34-kDa proteins were purified by electroelution (Figure 1A). To assess the vaccine potential of these antigens, we encapsulated them in cationic DSPC liposomes and administered them to BALB/c mice, followed by challenge with the *L. donovani* AG83 strain. Infection with this strain in BALB/c mice leads to progressive liver and spleen infection, resulting in hepatomegaly and splenomegaly within 3-4 months (27, 28). As previously observed, liposomal gp63 vaccination via the intraperitoneal (i.p.) route provided protection, with mice showing reduced parasite burden when sacrificed 3 months post-infection. In contrast, liposomal 34-kDa vaccination did not confer protection after the same duration of infection (Figure 1B and 1C). Control mice injected with PBS or empty liposomes exhibited high parasite loads, indicating a progressive infection.

**FIG 1.**
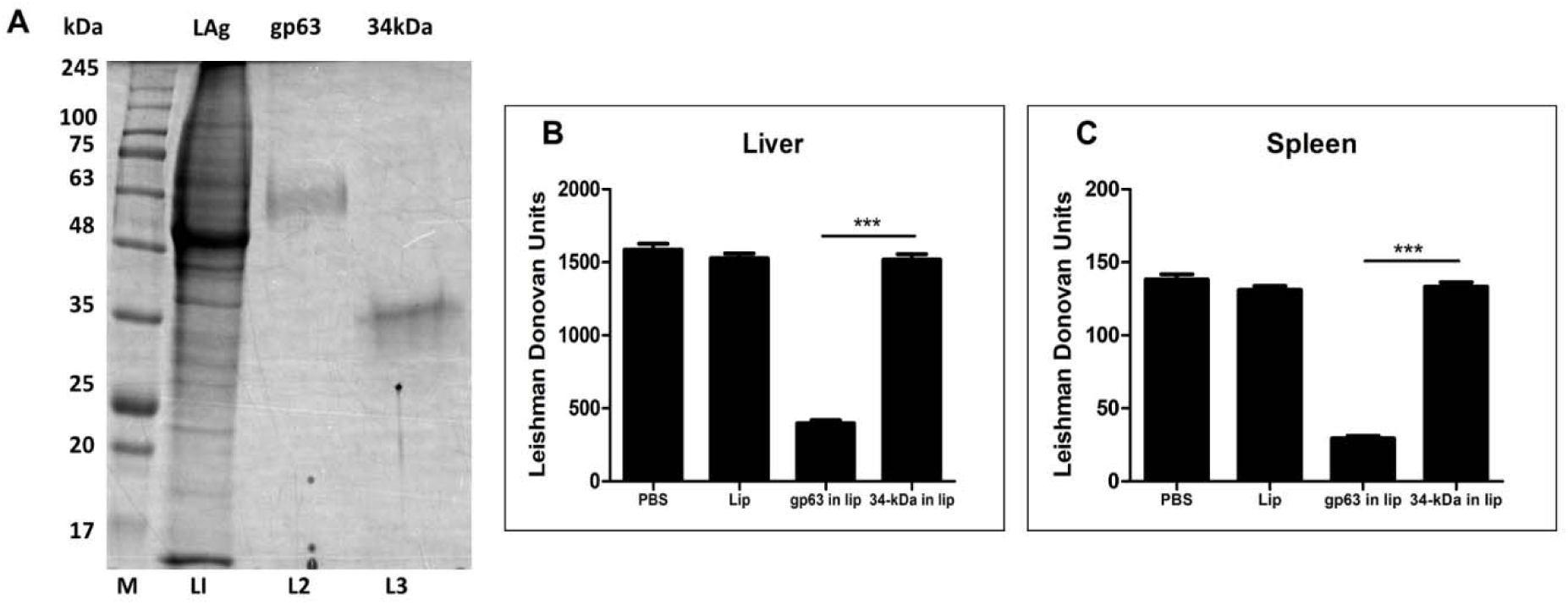
Characterization and clinical outcome of immunization against *L. donovani*. (A) SDS- PAGE analysis of electroeluted proteins, including leishmanial membrane antigens (LAg) (lane 1), gp63 (lane 2), and 34-kDa protein (lane 3), stained with Coomassie Blue. Molecular mass markers (left) are indicated in kilodaltons. Clinical outcome following *L. donovani* challenge in immunized BALB/c mice. Mice were vaccinated intraperitoneally three times at 14-day intervals with gp63 and 34-kDa protein entrapped in liposomes (2.5 µg/dose/animal), while control groups received PBS. Ten days post-immunization, the mice were challenged with 2.5 × 10^7^ promastigotes of *L. donovani*. Liver (B) and spleen (C) parasite burdens were measured 3 months post-infection as Leishman Donovan Units (LDU). Results are presented as mean ± S.E. (n=5). *** *P* < 0.001 compared to gp63 in liposome group.

### Cellular responses induced by liposomal gp63, but not by liposomal LACK vaccination

To assess the cellular immune responses, we evaluated delayed-type hypersensitivity (DTH) and splenocyte proliferation following footpad injection with LAg or in vitro stimulation with specific antigens, 10 days after the final vaccination. Mice vaccinated with liposomal gp63 exhibited a robust DTH response, while mice receiving liposomal 34-kDa vaccination did not show a significant response (Figure 2A). Similarly, splenocytes from mice immunized with liposomal gp63 displayed a marked proliferative response, whereas splenocytes from 34-kDa vaccinated mice showed no such response (Figure 2B). These results indicate that, despite being encapsulated in cationic DSPC liposomes, the 34-kDa antigen failed to induce a cellular immune response in BALB/c mice following vaccination.

**FIG 2.**
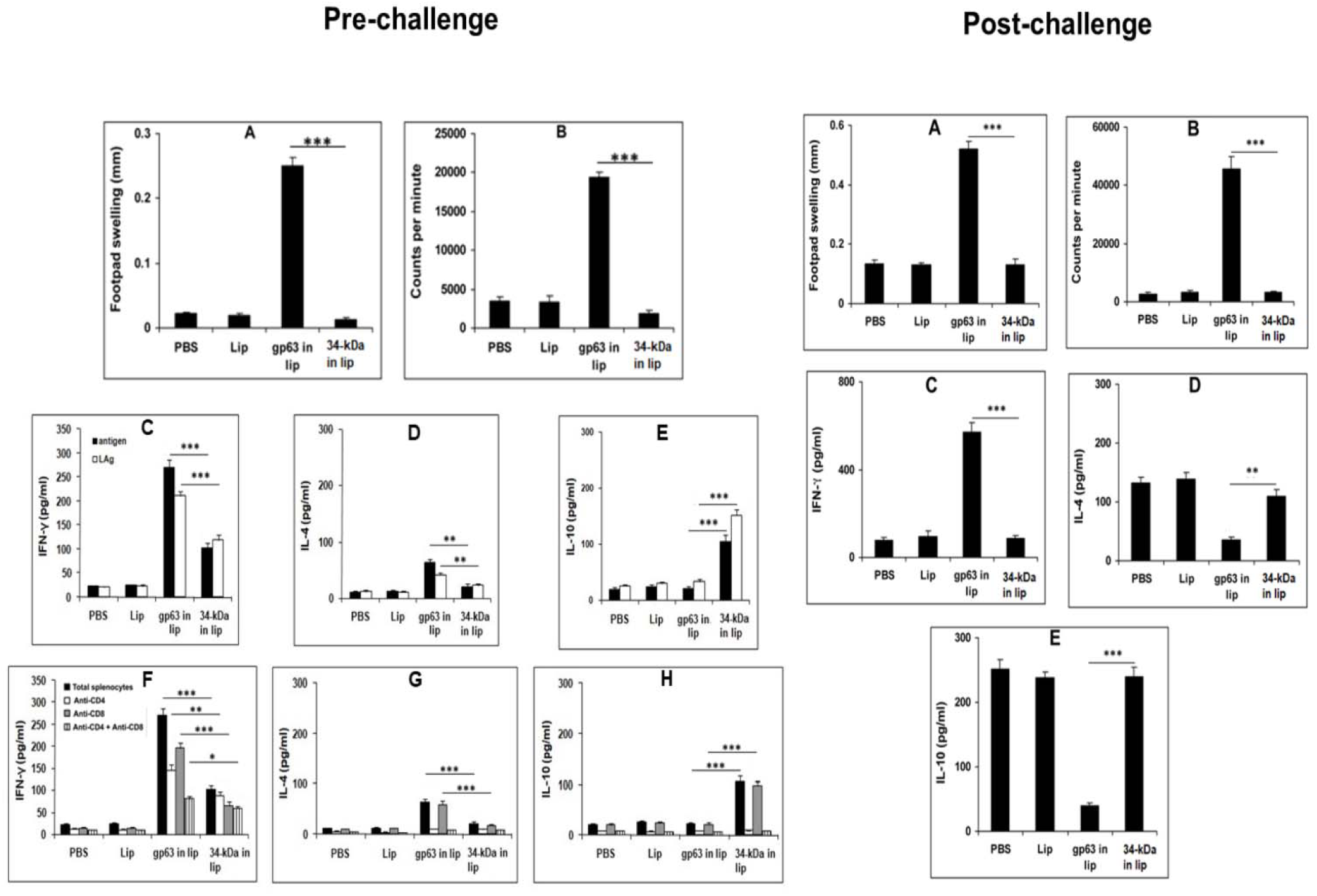
Pre- and post-challenge immunological responses in mice vaccinated with liposomal electroeluted antigens. Pre-challenge: (A) DTH response measured as the difference (in mm) between the thickness of test (40 µg LAg-injected) and control (PBS-injected) footpads after 24 h in mice immunized twice at 14-day intervals. (B) Proliferation of splenocytes from vaccinated mice, stimulated with LAg (10 μg/ml) for 96 hrs, pulsed with 1 μCi of [3H]-thymidine per well 18 h before harvesting on glass fiber paper. Thymidine uptake was measured in a β-scintillation counter. Post-immunization cytokine analysis: splenocytes from vaccinated and control mice were stimulated with LAg (10 μg/ml) and gp63/34-kDa (2.5 μg/ml) in the presence or absence of anti-CD4 and anti-CD8 antibodies 10 days after the last immunization. IFN-γ (C, F), IL-4 (D, G), and IL-10 (E, H) concentrations in the culture supernatants were quantified by ELISA, with each sample examined in duplicate. Post-challenge: (A) DTH response measured similarly as above. (B) Proliferation of splenocytes from infected mice stimulated with LAg (10 μg/ml) and analyzed for thymidine uptake as described. (C) Levels of IFN-γ (C), IL-4 (D), and IL-10 (E) in culture supernatants from splenocytes of infected mice stimulated with gp63/34-kDa (2.5 μg/ml) were quantified by ELISA. All results represent mean ± S.E. (n=5). * *P* < 0.05, ** *P* < 0.01 and *** *P* < 0.001 in comparison to gp63 in liposome group.

### Vaccination with liposomal gp63 and LACK induced differential IFN-**γ**, IL-4 and IL-10 production

IFN-γ, IL-4, and IL-10 are key cytokines known to play critical roles in determining the clinical outcome of infections. Therefore, we measured the levels of these cytokines produced by splenocytes from mice after in vitro restimulation with LAg or specific antigens. Splenocytes from mice immunized with liposomal gp63 exhibited higher levels of both IFN-γ and IL-4 (Figure 2C and 2D, Pre-challenge). In contrast, liposomal 34-kDa vaccination induced lower, though significant, levels of IFN-γ but no detectable IL-4 production. Notably, splenocytes from liposomal 34-kDa immunized mice released significant amounts of IL-10 when stimulated with LAg and specific antigens, whereas liposomal gp63 vaccination did not induce IL-10 production (Figure 2E).

To assess the relative contributions of CD4+ and CD8+ T cells in the production of these cytokines, blocking experiments were performed using anti-CD4 and anti-CD8 antibodies. The addition of both anti-CD4 and anti-CD8 antibodies to cultures from liposomal gp63 immunized mice inhibited the production of IFN-γ, indicating that both CD4+ and CD8+ T cells contribute to IFN-γ production (Figure 2F). In contrast, IL-4 production was solely attributed to CD4+ T cells, as demonstrated by blocking experiments (Figure 2G). For liposomal LACK immunized mice, IFN-γ production was observed from non-T cells, while IL-10 was primarily released by CD4+ T cells in the culture (Figure 2H).

### Challenge infection further polarizes IFN-**γ** production in mice vaccinated with liposomal gp63 and IL-10 in liposomal 34-kDa immunized mice

Three months post-challenge with *L. donovani* AG83, we reassessed the delayed-type hypersensitivity (DTH) response. Mice vaccinated with liposomal gp63 exhibited a robust DTH response, while those vaccinated with liposomal 34-kDa failed to show any response (Figure 2A, Post-challenge). We also evaluated the T cell proliferation response of immunized mice after challenge. Liposomal gp63 vaccination induced significant T cell proliferation, correlating with the strong DTH response. In contrast, liposomal 34-kDa did not elicit a similar response (Figure 2B). To assess cytokine production, we measured levels of antigen-specific IFN-γ, IL-4, and IL- 10 in culture supernatants of splenocytes isolated from vaccinated mice after 3 months of infection. Mice vaccinated with liposomal gp63 showed elevated levels of IFN-γ, a Th1 cytokine, indicating a robust Th1 response. Control mice, infected with *L. donovani* AG83, displayed high IL-4 levels (Figure 2D), consistent with previous findings in BALB/c mice (29). While liposomal gp63 vaccination reduced IL-4 levels compared to PBS and empty liposome controls, liposomal 34-kDa immunized mice had IL-4 levels similar to those of control groups, suggesting a lack of protection. Furthermore, mice vaccinated with liposomal 34-kDa had significantly higher levels of IL-10, whereas liposomal gp63 vaccination resulted in lower IL-10 production (Figure 2E). Elevated levels of IL-4 and IL-10 in the liposomal 34-kDa group may contribute to the failure of protection observed in these mice.

### Cloning, overexpression and purification of LACK from *L. donovani*

For a more precise evaluation of this electroeluted protein as a vaccine candidate, found to be LACK (30), we cloned and overexpressed the protein in recombinant form. *L. donovani* promastigotes (3rd-day cultures from the 2nd passage) were pelleted, and their genomic DNA was isolated using a genomic DNA isolation kit. The purity of the isolated DNA was confirmed by agarose gel electrophoresis (Figure 3A). The LACK gene was successfully amplified from *L. donovani* AG83 genomic DNA, as confirmed by a prominent 1 kb band (Figure 3B). The PCR products were purified and digested with NcoI and HindIII, and the pET28a vector was similarly digested. The vector and insert were ligated, transformed into DH5-α competent cells, and the resulting colonies were screened for correct ligation by plasmid digestion (Figure 3C). Positive clones were selected and used for the expression of LACK protein in the Rosetta strain of *E. coli*.

**FIG 3.**
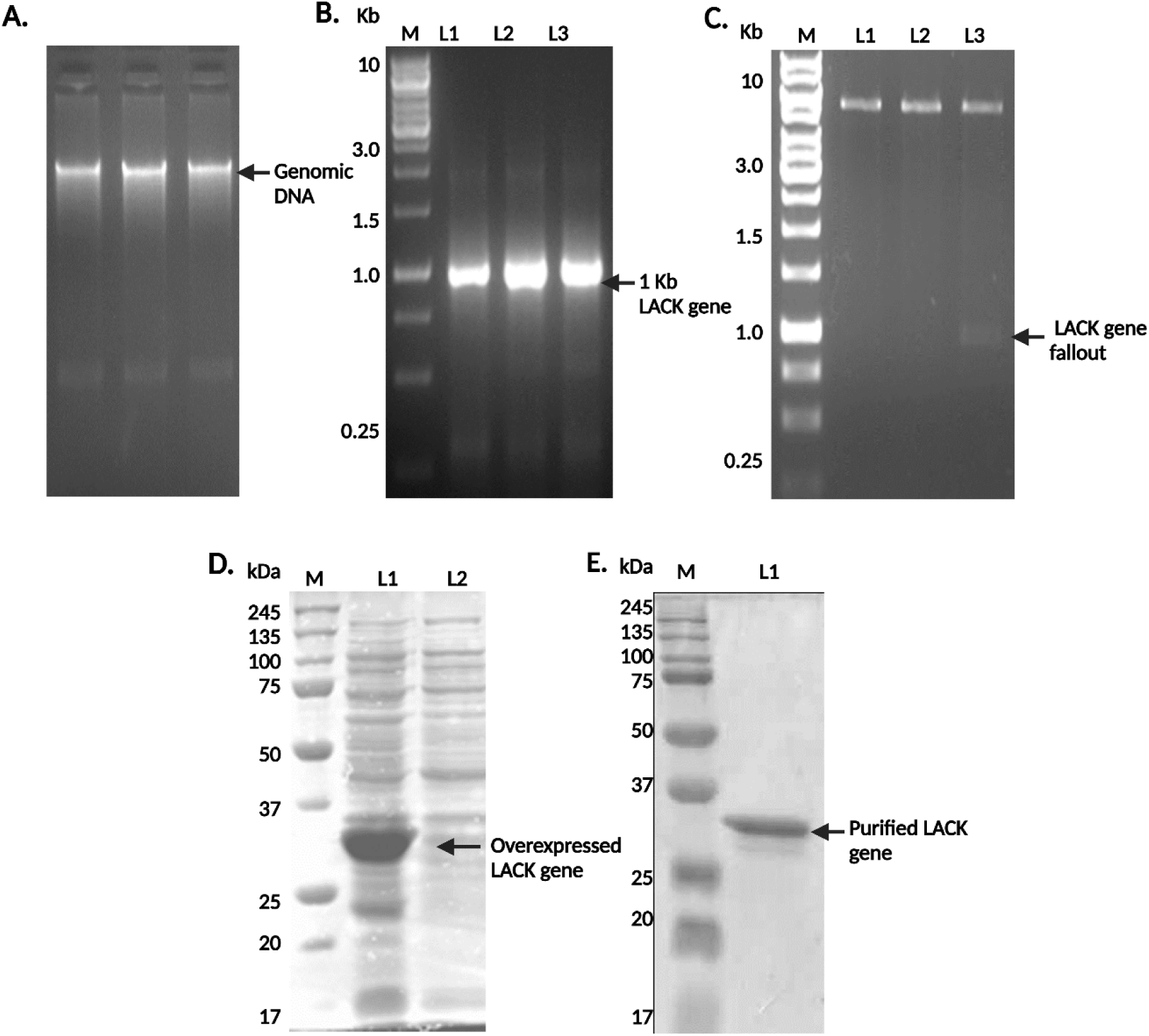
Cloning, Expression, and Purification of Recombinant LACK Protein from *L. donovani*. (A) Genomic DNA of *L. donovani* was isolated and used to amplify the LACK gene. (B) The amplified LACK gene was cloned into the pET28a vector and double-digested with NcoI and HindIII, yielding a 1 kb insert marked as L3. (C) Expression of the cloned LACK gene was confirmed in the Rosetta strain of *E. coli*, with protein induction marked as L1 compared to the uninduced control L2. (D) The recombinant LACK protein was purified and visualized by Coomassie staining.

The recombinant protein was overexpressed following induction with IPTG, and bacterial pellets were collected. SDS-PAGE analysis showed a thick band near 34 kDa, confirming the successful overexpression of LACK (Figure 3D). The protein was then purified using Ni-NTA chromatography, and its purity was verified by SDS-PAGE (Figure 3E). The purified LACK protein was subsequently entrapped in cationic DSPC liposomes for use in vaccination studies.

### Immunization of BALB/c mice with liposomal LACK leads to a skewed immune response favoring Th2 activation

To assess whether recombinant LACK could improve the immune responses induced by electroeluted 34-kDa protein, BALB/c mice were immunized intraperitoneally (i.p.) three times with free LACK, liposomal LACK, PBS, or empty liposome as controls. Ten days after the final immunization, DTH, antibody, and cytokine responses were evaluated. For the DTH response, swelling in the footpads of mice injected with LACK antigen and PBS was measured. Liposomal LACK-immunized mice showed no significant increase in antigen-specific DTH response compared to the control groups (Figure 4A, Pre-challenge), suggesting an absence of a cell-mediated immune response induced by LACK. Regarding antibody responses, sera from immunized mice were analyzed for LACK-specific antibodies. Immunized animals exhibited significantly higher antibody titers compared to controls (Figure 4B), indicating that LACK immunization induces humoral responses. Levels of IgG2a, associated with Th1 responses, and IgG1, associated with Th2 responses, were also measured. Liposomal LACK vaccination led to high titers of both IgG2a and IgG1 (Figure 4C, 4D). The IgG1:IgG2a ratio (Figure 4E) revealed a Th2-biased immune response following liposomal LACK vaccination.

**FIG 4.**
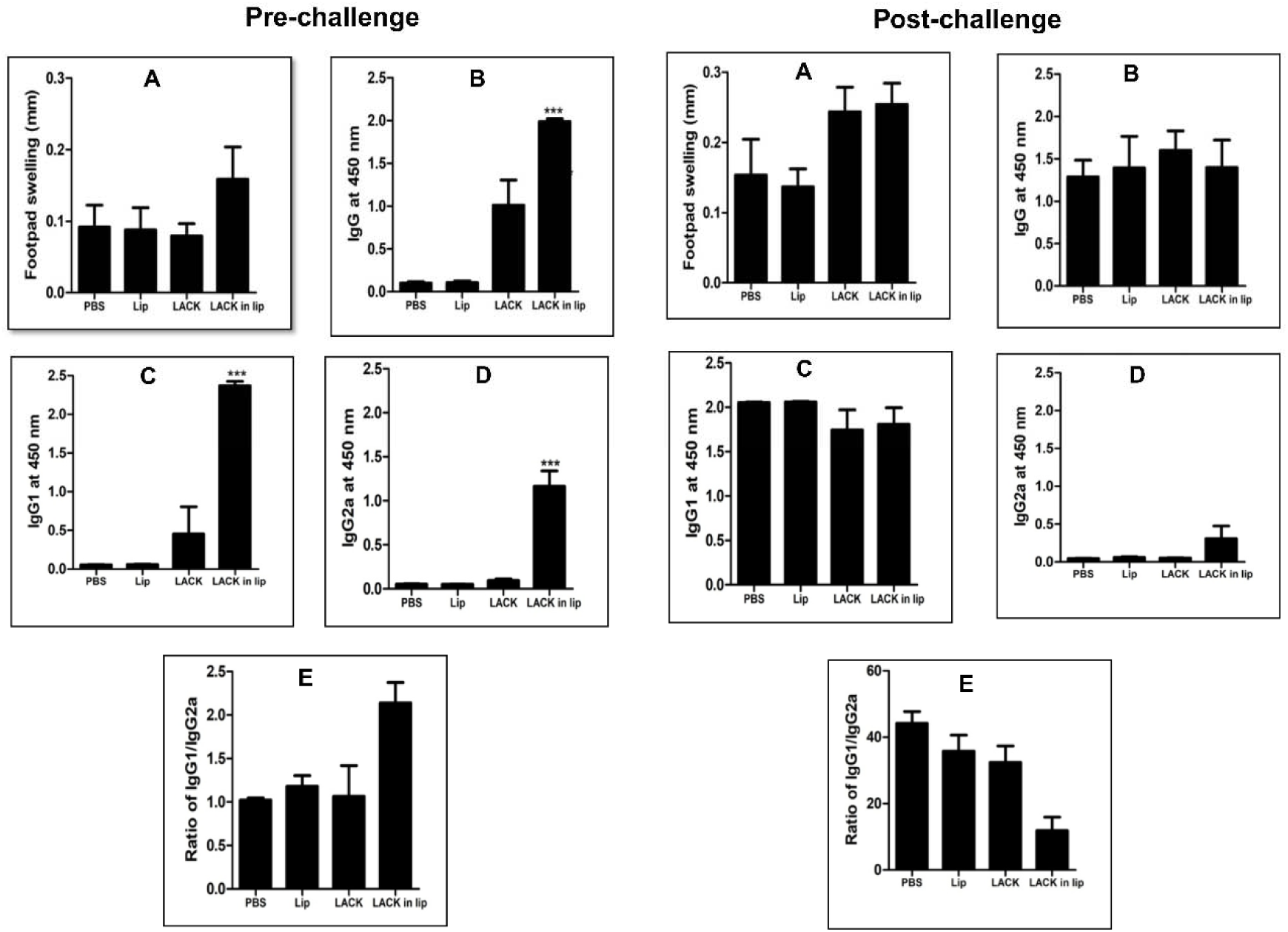
Pre- and post-challenge DTH and antibody responses in mice vaccinated with LACK alone or in liposomal formulations. Pre-challenge: Mice were immunized with three intraperitoneal injections of recombinant LACK either free in PBS or formulated in cationic liposomes at 14-day intervals. (A) DTH response was measured as the difference (in mm) between the thickness of the test (2.5 µg LACK-injected) and control (PBS-injected) footpads after 24 h. Serum samples collected 10 days after the last booster dose were assayed for LACK- specific IgG (B), IgG1 (C), and IgG2a (D) antibodies by ELISA at a dilution of 1:2000, with the ratio of IgG1/IgG2a (E) plotted. Post-challenge: Following a challenge with 2.5 × 10^7^ promastigotes of *L. donovani* 10 days post-immunization, DTH response (A) and serum antibody levels (B-E) were assessed 3 months post-infection. Results are presented as above, indicating the effects of vaccination formulation on immunological responses. All results represent mean ± S.E. (n=4), *** *P* < 0.001 in comparison to control groups.

Ten days after the final immunization, mice were challenged with *L. donovani* promastigotes. Ninety days post-challenge, DTH and antibody responses were reassessed. As with pre-challenge results, liposomal LACK-immunized mice showed no significant increase in DTH compared to control groups, indicating a lack of cell-mediated immune response (Figure 4A, Post-challenge). Antibody analysis revealed that all infected mice produced IgG antibodies against LACK (Figure 4B), with a uniform increase in IgG1, indicating a predominant Th2 response across all groups, including liposomal LACK-immunized mice, as evidenced by the IgG1:IgG2a ratio (Figure 4E).

### Immunization with LACK fails to down-regulate IL-4, IL-10 and TGF-**β** in immunized and mice challenged with *L. donovani*

To determine the specific immune response induced, splenocytes from immunized mice were antigen-pulsed with LACK, and cytokine levels in the culture supernatants were measured. Significant elevations in IL-4, IL-10, and TGF-β were observed in both LACK and liposomal LACK-immunized mice (Figure 5C, 5D, 5E, Pre-challenge). No significant increase in IL-12 was noted in the vaccinated groups compared to controls (Figure 5B), although both LACK and liposomal LACK immunization resulted in significantly higher levels of IFN-γ (Figure 5A) compared to controls.

**FIG 5.**
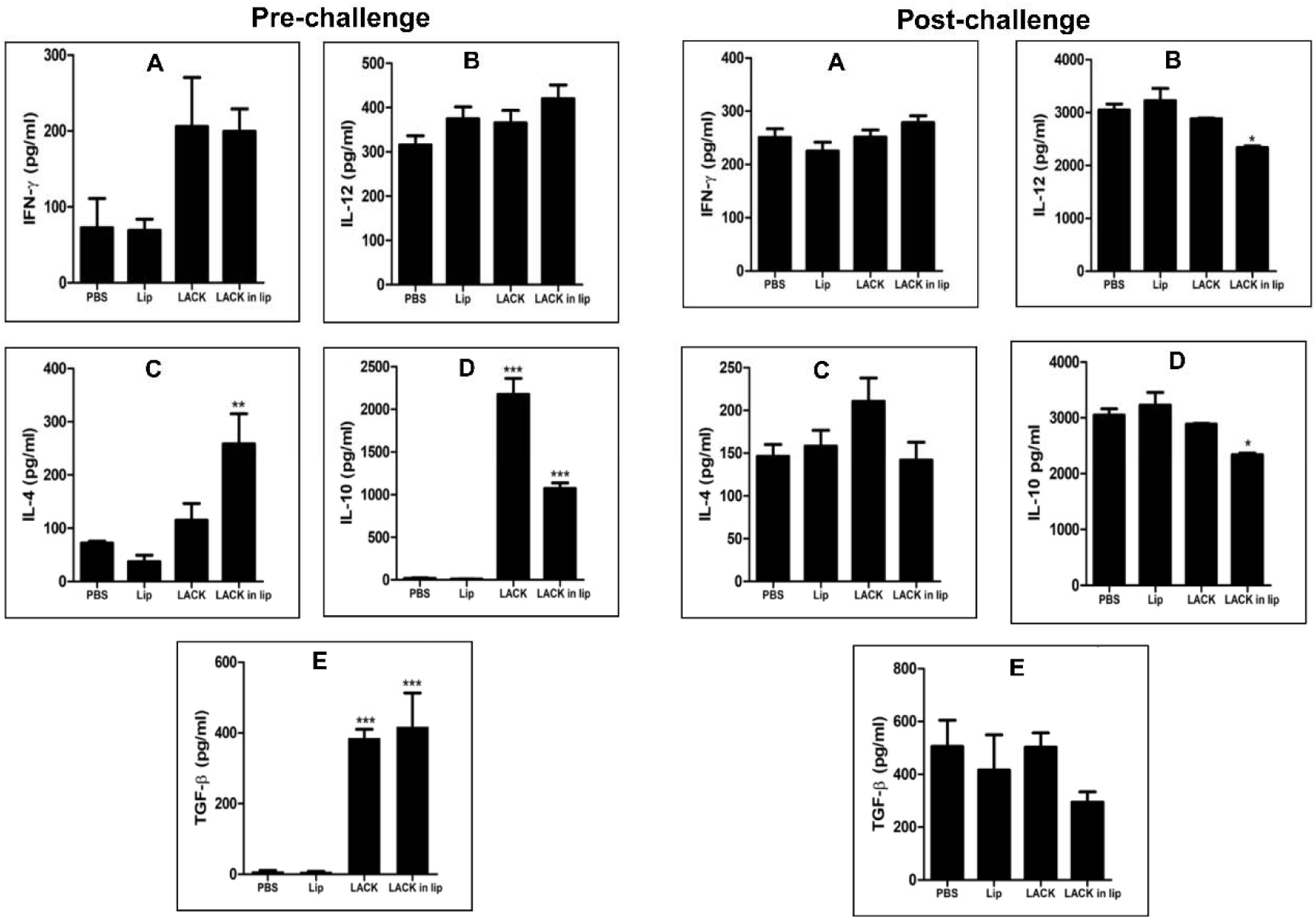
Pre- and post-challenge cytokine responses in mice immunized with LACK alone or in liposomal formulation. Pre-challenge: Mice received three intraperitoneal injections of LACK, either free in PBS or formulated in cationic liposomes, at 14-day intervals (2.5 µg/dose/animal). (A) Ten days after the last immunization, splenocytes were cultured and stimulated with LACK (2.5 µg/ml) for 72 hr, with concentrations of released IFN-γ (A), IL-12 (B), IL-4 (C), IL-10 (D), and TGF-β (E) in the culture supernatants quantified by ELISA. Post-challenge: Following a challenge with 2.5 × 10^7^ promastigotes of *L. donovani* 10 days post-immunization, splenocytes were again cultured and stimulated with LACK (2.5 µg/ml) 3 months post-infection. Cytokine levels (A) IFN-γ, (B) IL-12, (C) IL-4, (D) IL-10, and (E) TGF-β were measured in culture supernatants using ELISA. All results are presented as mean ± S.E. (n=4), * *P* < 0.05 in comparison to control groups, highlighting the impact of vaccination formulation on cytokine responses \**P* < 0.05, ** *P* < 0.01, *** *P* < 0.001 in comparison to control groups.

In order to determine the immunomodulation following challenge infection with *L. donovani*, mice were challenged with virulent promastigotes ten days after the last vaccination, and at ninety days after infection the animals were sacrificed and the splenocytes were stimulated with LACK. The cytokines in the culture supernatants were estimated after 72 hrs of stimulation.

After challenge infection with *L. donovani*, mice were sacrificed at 90 days post-infection, and their splenocytes were re-stimulated with LACK. Cytokine levels in the culture supernatants were measured after 72 hours of stimulation. Post-challenge, all groups exhibited high levels of IL-4, IL-10, and TGF-β (Figure 5C, 5D, 5E), though there was some downregulation of IL-10 and TGF-β in the liposomal LACK-vaccinated group. Interestingly, while IL-12 levels (Figure 5B) slightly decreased in liposomal LACK-vaccinated mice, there was no change in IFN-γ levels (Figure 5A) across all groups.

### Immunization with liposomal LACK failed to protect BALB/c mice from *L. donovani* infection

To assess the correlation between vaccine-induced immune modulation and protective efficacy, the parasite burden in the liver and spleen of infected mice was determined. The Giemsa-stained smears of liver and spleen samples from control and vaccinated mice were microscopically examined to quantify the parasite load (LDU). Mice immunized with liposomal LACK exhibited no significant reduction in parasite load in the liver (Figure 6A) and only a slight decrease in the spleen (Figure 6B) compared to PBS-immunized controls. Additionally, liver and spleen samples were further analyzed using the Limiting Dilution Assay (LDA) to assess the presence of live parasites. The results showed comparable titration of parasites between both control and immunized mice, indicating that LACK and liposomal LACK failed to protect against *L. donovani* infection (Figure 6C and 6D).

**FIG 6.**
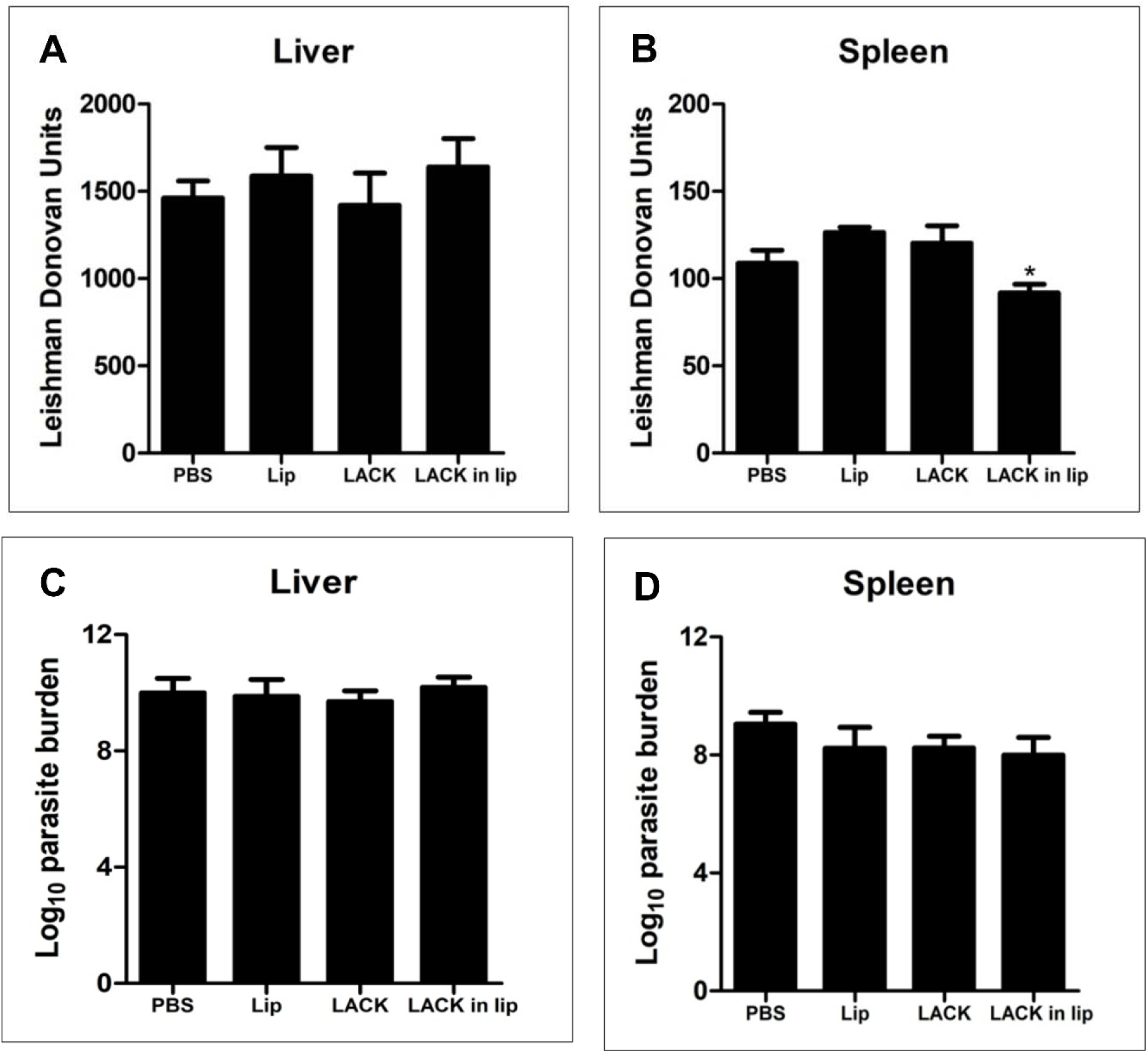
Clinical outcome following *L. donovani* challenge in BALB/c mice immunized with LACK. Mice were vaccinated intraperitoneally three times with 14 days interval with free LACK or entrapped in liposomes (2.5 µg/dose/animal). Control groups received only PBS. Ten days post-immunization, the mice were challenged with 2.5 × 10^7^ promastigotes of *L. donovani.* Liver (A, C) and spleen (B, D) parasite burdens were measured 3 months post infection as Leishman Donovan Units (LDU) (A,B), and log_10_ parasite burden (C, D), respectively. Results represent mean ± S.E. (n=5), * *P* < 0.05 in comparison to control groups.

### Cytokine producing CD4 and CD8 T cells from splenocytes of immunized and post infection mice

To identify the cellular source of the cytokine response, splenocytes from immunized and infected mice were analyzed by flow cytometry. Stimulated splenocytes were stained with anti- CD3-APC Cy7, anti-CD4-BV50, and anti-CD8 Per CP Cy 5.5 antibodies, followed by intracellular staining for cytokines using anti-IFN-γ-BV421, anti-IL-12-APC, anti-IL-2-R718, anti-IL-4-PECF594, and anti-IL-10-PE antibodies. The gating strategy for surface antigens (CD3, CD4, CD8) and intracellular cytokines (IFN-γ, IL-12, IL-2, IL-4, IL-10) is shown in Figure 7A and Figure 8A.

**FIG 7.**
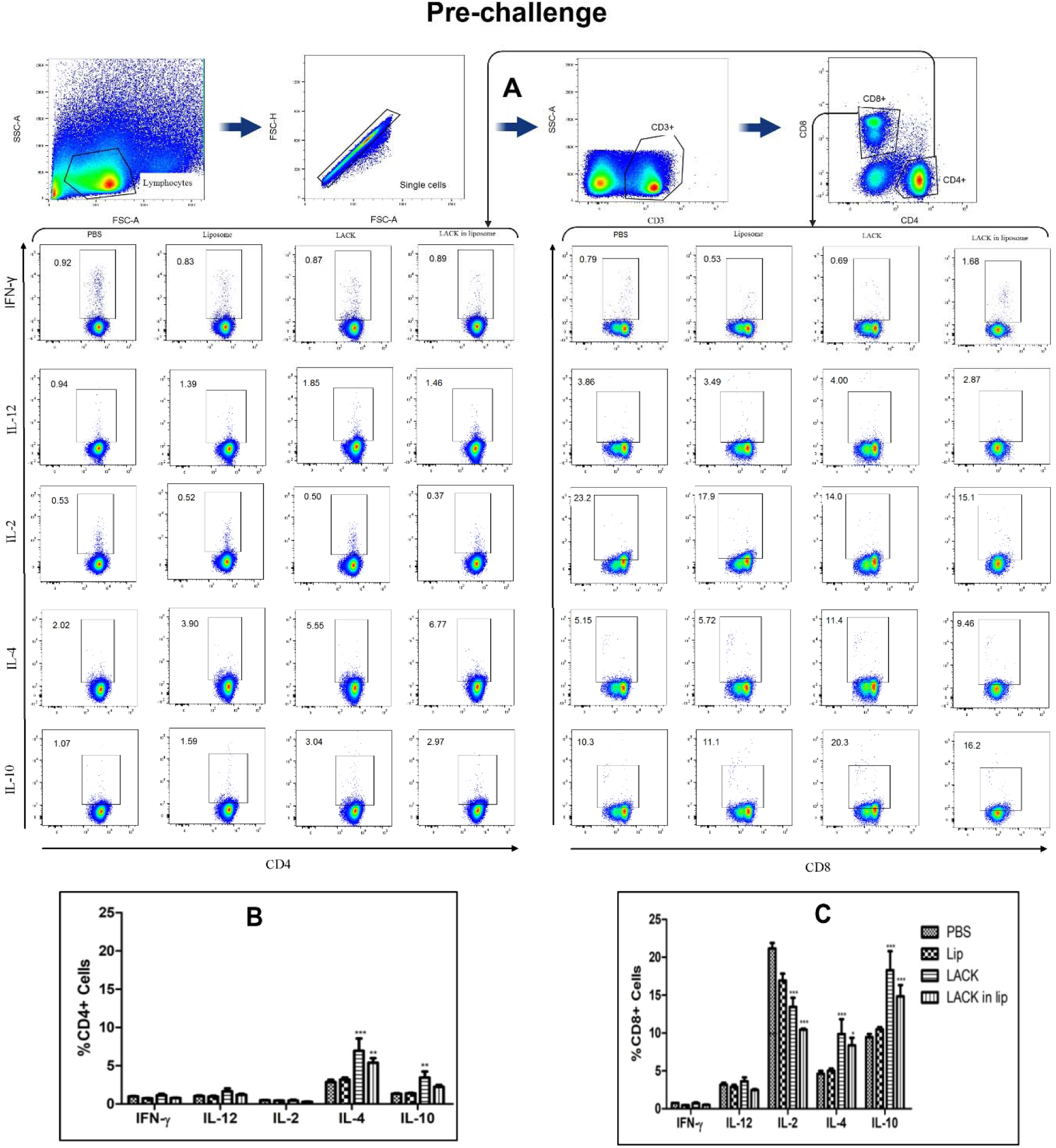
LACK induced T cell response in immunized BALB/c mice. To determine the frequencies of antigen-specific post immunization cytokine producing CD4+ and CD8+ T cells, LACK stimulated splenocytes were isolated from differently immunized groups of mice and stained for surface markers and different cytokines. Cells were sorted by multiparameteric flow cytometry and analyzed with FlowJo software. (A) Representative plots showing frequencies of IFN-γ, IL-12, IL-2, IL-4 and IL-10 producing CD4+ and CD8+T cells. Frequencies of CD4^+^ (B) and CD8^+^ (C) T cells expressing IFN-γ, IL-12, IL-2, IL-4 and IL-10. Results represent mean ± S.E. (n=4), \**P* < 0.05, ** *P* < 0.01, *** *P* < 0.001 in comparison to control groups.

**FIG 8.**
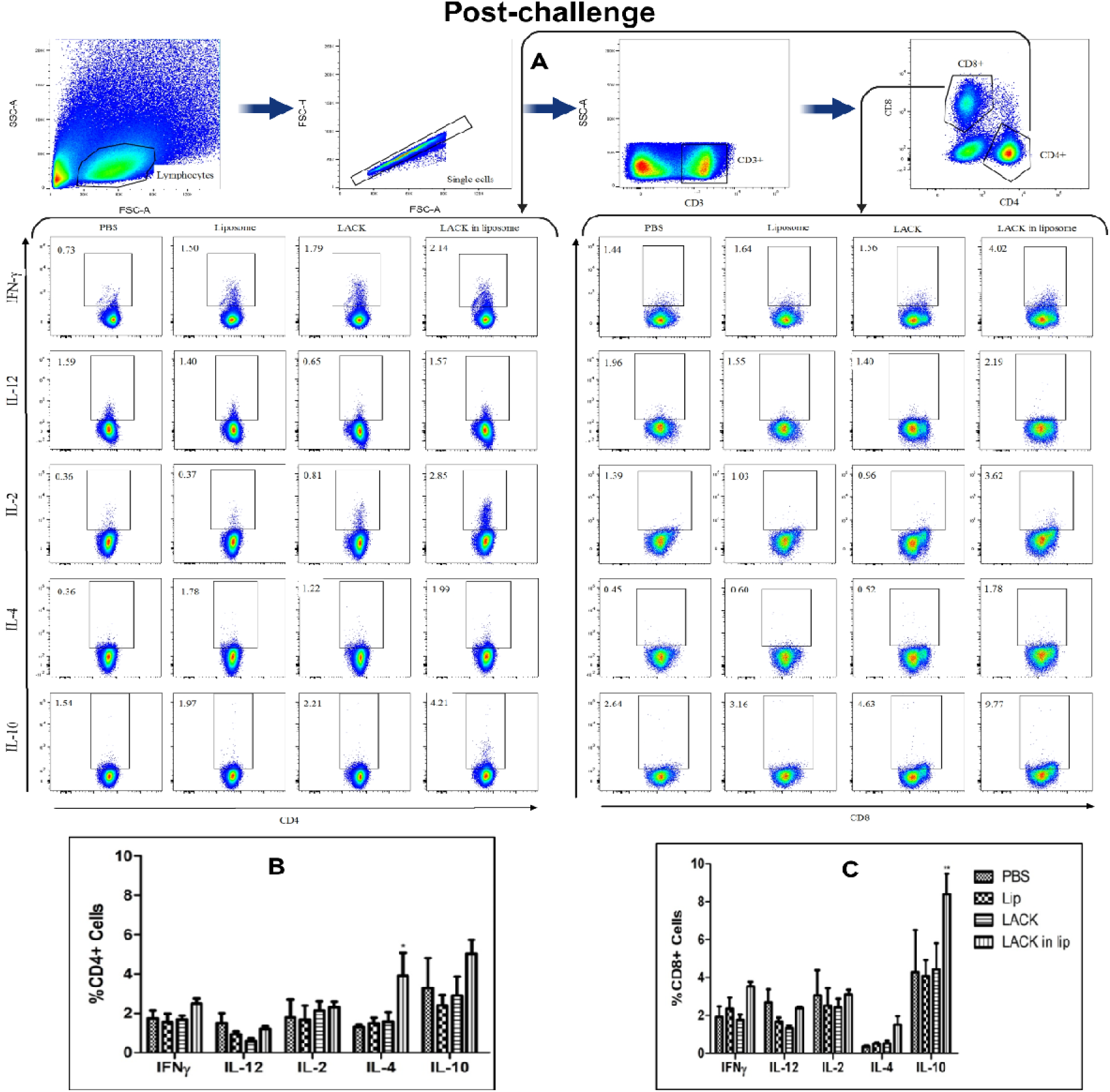
LACK induced T cell response in immunized BALB/c mice infected with *L. donovani*. To determine the frequencies of antigen-specific post infection cytokine producing CD4+ and CD8+ T cells, LACK stimulated splenocytes isolated from differently immunized groups of mice and stained for various surface markers and different cytokines. Cells were sorted by multiparameteric flow cytometry and analyses with FlowJo software to determine the frequencies of T cells. (A) Representative plots showing total frequencies of IFN-γ, IL-12, IL-2, IL-4 and IL-10 producing CD4+ and CD8+ T cells. Frequencies of CD4+ (B) and CD8+ (C) T cells expressing IFN-γ, IL-12, IL-2, IL-4 and IL-10. Results represent mean ± S.E. (n=5). * *P* < 0.05, ** *P* < 0.01 in comparison to control groups.

In vaccinated mice, the gated CD4+ T cells exhibited an increased proportion of IL-4- and IL-10- producing cells (Figure 7B). CD8+ T cells from liposomal LACK-immunized mice also showed higher proportions of IL-4 and IL-10 producers and fewer IL-2-producing cells compared to CD8+ T cells from PBS-immunized mice (Figure 7). After infection, the proportions of CD4+ and CD8+ T cells producing intracellular cytokines remained largely unchanged across all groups. A slight increase in IL-4-producing CD4+ T cells (Figure 8B) and IL-10-producing CD8+ T cells (Figure 8C) was observed in the liposomal LACK group, but no significant protective immune response was detected, further indicating the lack of efficacy of LACK and liposomal LACK in providing protection against *L. donovani* infection.

## DISCUSSION

LACK, a highly conserved antigen among *Leishmania* species, has been proposed as a potential vaccine candidate for both cutaneous and visceral leishmaniasis (31, 32). In previous studies, we reported on the efficacy of LAg in association with cationic liposomes, with DSPC-bearing vesicles demonstrating the most significant protective effect (33). The rationale for reassessing LACK in this study was grounded in its consistent recognition by sera from kala-azar patients and its inclusion in the LAg pool, which comprises proteins ranging from 18 to 190 kDa, including gp63 and a 34-kDa component identified as LACK (30, 34). While earlier studies using DNA or peptide-based LACK formulations yielded suboptimal protection, we hypothesized that this might be attributed to insufficient delivery and immune potentiation rather than intrinsic antigenic limitations. Therefore, we employed DSPC-based cationic liposomes, a delivery system validated by previous studies (24–26, 35) to assess whether the optimized delivery platform could enhance LACK’s immunogenic potential.

In this comparative study, we evaluated the immune responses and challenge outcomes in BALB/c mice immunized with either gp63 or LACK formulated in DSPC-bearing cationic liposomes. Both vaccine formulations elicited IL-4 and IFN-γ responses, but cytokine levels were consistently lower in the LACK immunised group. Importantly, high levels of IL-10 detected prior to challenge in LACK-immunized mice were closely associated with vaccine failure. In contrast, gp63-immunized mice exhibited minimal IL-10 production and achieved significant protection. Furthermore, elevated TGF-β levels in the LACK group prior to challenge underscored the roles of IL-10 and TGF-β as negative correlates of protection in this murine VL model.

The adaptive immune system, comprising T cells and B cells, mediates cell-mediated immunity (CMI) and humoral immunity, respectively. CMI is particularly crucial for protection against intracellular pathogens like *Leishmania* (36), and delayed-type hypersensitivity (DTH) is often used as an indicator of CMI activation in vivo (37). To assess CMI, DTH response was measured. Our findings revealed that gp63 delivered using cationic liposomes effectively induced a strong DTH response, indicating a robust CMI, consistent with previous studies that highlighted the immunogenicity of gp63 as a vaccine candidate (26). In contrast, LACK, even when delivered in the same cationic liposome formulation, showed limited ability to elicit CMI. We next evaluated the humoral response, using gp63, known for its strong IgG response, as a benchmark (35). Both LACK and its liposomal formulation induced comparable robust IgG responses, demonstrating their immunogenicity. The kinetics of IgG2a and IgG1 production, which reflect Th1 and Th2 responses respectively, were also measured (38). Earlier studies on gp63 showed a Th1-driven response, marked by higher IgG2a and lower IgG1 levels, resulting in a high IgG2a/IgG1 ratio(35). In contrast, our study showed that immunization with liposomal LACK resulted in upregulation of IgG1 and a low expression of IgG2a, leading to a high IgG1/IgG2a ratio, indicative of a Th2-driven immune response.

To further evaluate the CMI response elicited by CD4+ T cell subsets in response to the two different antigens presented in liposomes, we conducted cytokine analysis of splenocytes from immunized mice. In the context of visceral leishmaniasis (VL), the balance between Th1 and Th2 responses plays a critical role in determining disease outcomes. Th1 responses, characterized by cytokines such as IFN-γ, are crucial for controlling intracellular pathogens like *Leishmania*, as they activate macrophages and promote pathogen clearance. In contrast, Th2 responses, marked by cytokines like IL-4, IL-10, and TGF-β, are associated with disease progression and parasite persistence, as they suppress effective macrophage activation and promote an anti-inflammatory response (39). Both gp63 and LACK immunization induced production of IFN-γ and IL-4. However, the levels of these cytokines were significantly lower in the LACK-vaccinated group. Notably, LACK immunization induced higher levels of Th2 cytokines, particularly IL-10 and TGF-β, a pattern not observed in the gp63-vaccinated mice, which showed lower levels of these cytokines. This suggests that while both antigens stimulate an immune response, LACK induces a Th2-dominated response (40), whereas gp63 primarily drives a Th1-biased response (26, 34). Following challenge with virulent *L. donovani* promastigotes, LACK immunization failed to downregulate the anti-inflammatory cytokines IL- 4, IL-10, and TGF-β, potentially leading to disease progression. In contrast, gp63 immunization, which promoted the production of IFN-γ and downregulation of IL-10 and IL-4 was able to control the infection. This was reflected in the significantly lower parasite load observed in the gp63-vaccinated group compared to the LACK and control groups, as determined by LDU and LDA. Elevated levels of IL-4, IL-10, and TGF-β in LACK-immunized mice correlated with higher parasite burdens in both the liver and spleen, while the Th1-driven response in gp63- immunized mice correlated with disease control (41).

To identify the antigen-specific cellular sources of cytokines, we performed ex vivo T cell depletion studies and flow cytometric (FACS) analysis (42). In the depletion study, we separately and simultaneously depleted CD4+ and CD8+ T cells from splenocytes to assess the impact on cytokine production by the remaining cells. This investigation aimed to elucidate the roles of CD4+ and CD8+ T cells in the immune response. Our results demonstrated that the cationic liposome formulation of gp63 effectively activated CD4+ and CD8+ T cells to produce IFN-γ, leading to protective effects in BALB/c mice challenged with *L. donovani*, consistent with previous reports (26). In contrast, in LACK-immunized mice, neither single nor combined depletion of CD4+ and CD8+ T cells resulted in any change in IFN-γ production. This suggests that CD4+ helper T cells and cytotoxic CD8+ T cells were not involved in the low-level IFN-γ production in LACK-vaccinated mice (43). which likely contributed to the failure to control parasite proliferation and disease progression (40). Multiparametric flow cytometric analysis with recombinant LACK further confirmed the involvement of antigen-specific CD4+ and CD8+ T cells. Pre-challenge data indicated that these T cell subsets were not responsible for the secretion of IFN-γ, albeit at low levels, suggesting that other cell populations might be responsible for its secretion. Instead, antigen-specific CD4+ and CD8+ T cells predominantly produced IL-4 and IL-10. The significant production of these cytokines in the LACK-vaccinated mice was associated with the failure of LACK to protect against *L. donovani*, even when administered with an optimized liposomal adjuvant system (44).

The immune response following immunization and the subsequent failure of LACK to protect against infection are intriguing. Despite both gp63 and LACK being encapsulated in DSPC- bearing cationic liposomes, only gp63 proved effective as a vaccine, while LACK failed. The failure of LACK even under these favorable delivery conditions suggests intrinsic limitations of the antigen itself and supports a reallocation of future vaccine efforts toward more promising targets. Nonetheless, reporting such negative outcomes is crucial for guiding rational vaccine design and preventing redundant efforts. Although potent liposomes are known to activate MHC pathways, including MHC I, and enhance the potency of subunit vaccines, LACK induced an anti-inflammatory immune response that was insufficient to confer protection against disease (45). LACK plays a significant role in the *Leishmania* lifecycle and infection process, making it a promising candidate for broad-spectrum vaccine development (46). However, studies have identified a dominant IL-10 epitope between amino acids 157 and 173 of LACK that induces IL- 10 production upon stimulation of PBMCs from cured and healthy humans (32). Consistent with these findings, we also observed that stimulation of VL-cured human PBMCs with LACK led to the highest production of IL-10 and TGF-β compared to other antigens (30). These results align with our immunization study, where we similarly observed upregulation of IL-10, TGF-β, and IL-4, contributing to a Th2-dominated response. The efficacy of any vaccine is influenced by multiple factors, and the failure of LACK as a vaccine candidate may be attributed to several reasons. While DNA vaccination can induce antibody responses, the magnitude of these responses is often lower than those generated by protein or live/attenuated vaccines (47). In a previous study, we demonstrated a dose-dependent protective efficacy of gp63 in liposomes, where increasing the dose initially enhanced CMI and improved disease protection. However, beyond a certain dose, further increases in the vaccine dose reduced protection, correlating with a decrease in CMI and an increase in the antibody response (26). In this study, the LACK dose (10 µg) was chosen based on the effective dose previously established for gp63 in our earlier studies. It is possible that a lower or higher dose of LACK, which could enhance CMI while minimizing antibody responses, might have been more effective to shift the Th2-biased response toward a more protective Th1 profile. Given the limitations of LACK as a standalone antigen, future strategies could involve its combination with more immunogenic antigens, such as gp63 or HASPB, to balance or override its Th2 polarization. Multivalent or cocktail vaccines incorporating multiple antigens could provide synergistic protection. Additionally, exploring alternative delivery systems or adjuvants such as MPLA, CpG-ODNs, virus-like particles (VLPs), or mRNA-based platforms may help redirect LACK-induced immunity toward a more desirable Th1-type response. While the BALB/c mouse model is a widely accepted preclinical model for VL vaccine research, it does not fully reflect the immunopathology of human VL. Therefore, future validation studies using complementary models such as Syrian golden hamsters or humanized mice would strengthen the generalizability and translational potential of these findings.

Cationic liposomes developed by our team typically exhibit a size range of 200-250 nm and a zeta potential of 50-60 mV, confirming their cationic nature (25). Consistent with this, our current formulations also showed similar characteristics, with liposomal LACK displaying a mean size of 215.46 ± 8.1 nm and a zeta potential of 51.8 ± 7.05 mV (Table 1). Lipids with high transition temperatures, such as DSPC, contribute to a stable lipid bilayer in cationic liposomes, facilitating uptake by antigen-presenting cells (APCs) through endocytosis (48). These characteristics are advantageous for MHC-mediated antigen presentation, leading to the generation of antigen-specific CD4+ and CD8+ T cell responses, which are essential for a Th1 response (49). While these responses have been successfully observed with liposomal gp63, they were not evident with LACK. Several studies have demonstrated that vesicle size significantly impacts the immune response. Specifically, an average size of 225 nm induces Th1 responses, characterized by increased IFN-γ and IgG2a, while a size of 155 nm elicits a Th2 response, marked by IgG1 titers and low or no IgG2a secretion (50, 51). Despite both gp63- and LACK- encapsulated cationic liposomes being within the size range of 250-300 nm, only the gp63 formulation demonstrated protective effects. The failure of liposomal LACK to induce a protective immune response may be due to antigen-dependent differences in cellular processing of LACK-associated cationic liposomes. Further investigations are needed to clarify these potential alterations and their impact on immune responses.

## MATERIALS AND METHODS

### Animals

BALB/c mice were used to evaluate the immunogenicity of the vaccine formulations. All animal studies were approved by the Institutional Animal Ethics Committee (IAEC) of CSIR-IICB and followed the guidelines of CPCSEA, Government of India.

### Maintenance of parasites

*L. donovani* strain AG83 (ATCC® PRA-413™) was maintained in hamsters at the animal housing facilities of CSIR-IICB, Kolkata. For parasite isolation, spleens from infected hamsters were aseptically removed, and amastigotes in the splenocytes were allowed to transform into promastigotes in Schneider’s insect medium containing 20% FBS, 100 µg/ml penicillin G, and streptomycin salts, incubated at 22°C for one week.

### Culture of *L. donovani* promastigotes

Freshly transformed *L. donovani* promastigotes were cultured in Medium 199 containing 10% FBS, 100 µg/ml penicillin G, and streptomycin salts at 22°C. The parasite density was maintained at 2×10 /ml by serial passaging. Parasites used for infection and experimental studies were between the 2nd and 4th passages.

### Preparation of antigens and Electroelution of proteins from SDS-PAGE gels

Stationary-phase *L. donovani* promastigotes were washed with cold PBS, resuspended in 5 mM Tris-HCl (pH 7.6), and vortexed. Following centrifugation, the membrane pellet was isolated, sonicated, and centrifuged again to obtain the supernatant containing leishmanial antigens (LAg). Protein concentration was determined using the Lowry method. The protein profile was analyzed by SDS-PAGE and visualized with Coomassie Brilliant Blue staining. Proteins (63 kDa and 34 kDa) were extracted from a 10% SDS-PAGE gel using an Electro-Eluter, dialyzed, lyophilized, and reconstituted in PBS (33).

### Cloning of LACK Gene

Genomic DNA of *L. donovani* AG83 (ATCC® PRA-413™) promastigotes was isolated, and the LACK gene was amplified using primers containing NcoI and HindIII restriction sites, with histidine codons added to the C-terminus. The PCR reaction was conducted under standard conditions, and the amplified fragment was cloned into the pET28a expression vector. The recombinant plasmid was confirmed by restriction digestion and agarose gel electrophoresis.

### Expression and Purification of LACK Protein

The plasmid containing the LACK gene was transformed into Rosetta strain *E. coli* for protein expression. Bacterial cultures were grown to log phase and induced with IPTG (0.5 mM) for 4 hours at 30°C. After harvesting, cells were lysed, and inclusion bodies were solubilized in 8M urea. Protein purification was performed using Ni-NTA agarose columns under denaturing conditions. The recombinant LACK protein was refolded by dialysis and stored in PBS.

### Entrapment of antigens in liposomes

Cationic liposomes were prepared using DSPC, cholesterol, and stearylamine (7:2:2 molar ratio) following previously described protocols (35). Lipids were dissolved in chloroform, and a thin film was formed via rotary evaporation under reduced pressure. The film was rehydrated with PBS containing either gp63, or 34-kDa antigen, or recombinant LACK, then vortexed and briefly sonicated. Antigen-loaded liposomes were separated from unentrapped proteins by ultracentrifugation. Protein content was measured using the Lowry method with 10% SDS. The size and surface charge of liposomes were analysed using dynamic light scattering (DLS) and zeta potential measurements with the Zetasizer Nano ZS (Malvern Instruments, U.K.).

### Immunization and infection schedule

BALB/c mice were immunized with gp63, 34-kDa antigen, or LACK in PBS or liposomes by intraperitoneal injection. Three injections of 2.5 µg protein were administered at two-week intervals in a total volume of 200 μl. Control groups received either PBS or empty liposomes. Ten days after the final booster, serum samples were collected, and spleens were harvested for immunological analyses. Another set of immunized animals was challenged via the tail vein with 2.5x10^7^ freshly transformed stationary phase promastigotes in 200 µl PBS and kept for 3 months (52).

### Cell proliferation

Spleen cells from immunized and infected mice were cultured in RPMI 1640 medium with 10% FBS. Red blood cells were lysed, and viable mononuclear cells were counted using Trypan blue exclusion. Cells were stimulated with gp63 or 34-kDa antigens in the presence or absence of anti-CD4 or anti-CD8 antibodies. Cell proliferation was measured by [3H]-thymidine incorporation, followed by counting using a β-scintillation counter (27).

### DTH

DTH responses were assessed in BALB/c mice ten days after the final immunization (third dose) administered at 14-day intervals. Mice received an injection of 40 µg LAg or 2.5 µg recombinant LACK protein into the right hind footpad, while the left hind footpad was injected with PBS as a control. After 24 hours, footpad swelling was measured using a digital caliper, and the difference in thickness between the antigen-injected and control footpads was recorded as the DTH response. To evaluate long-term recall responses, DTH measurements were also performed three months post-challenge with virulent *L. donovani* promastigotes (27, 53, 54).

### Determination of antibody response by ELISA

Serum samples from immunized and infected mice were analyzed for antigen-specific antibodies using ELISA. Microtiter plates were coated with gp63, 34-kDa antigen, or LACK (10 µg/ml), and sera were diluted 1:2000. HRP-conjugated goat anti-mouse IgG or isotype-specific antibodies (IgG1, IgG2a) were used for detection. Absorbance was measured at 450 nm (26).

### Cytokine ELISA

Spleen cells obtained from mice after immunization and 3 months after infection were RBC lysed and splenocytes were cultured in 12 well plates at the cellular concentration of 2x10^6^ cells/ well. The cultured splenocytes were stimulated with purified antigens at 2.5 µg/ml. Culture supernatants collected after 72 hrs were estimated for IFN-γ, IL-12, TGF-β, IL-4 and IL-10 by sandwich ELISA (BD Biosciences) (29).

### Identification and estimation of cytokine producing lymphocytes by flow cytometry

Splenocytes from immunized and infected mice were cultured and stimulated with recombinant LACK antigen. After 10–12 hours of stimulation, cells were stained for surface markers CD3, CD4, and CD8, followed by intracellular staining for cytokines including IFN-γ, IL-12, IL-2, IL- 4, and IL-10. Samples were acquired on a flow cytometer, and data were analyzed using a sequential gating strategy: lymphocyte population was first gated using FSC vs SSC, followed by exclusion of doublets (FSC-H vs FSC-A), gating on live cells, and subsequently CD3+ T cells were gated into CD4+ and CD8+ subsets. Intracellular cytokine-positive cells were then identified within each subset (24).

### LDU

Infection was evaluated by estimation of parasite burden in the liver and spleen in terms of Leishman-Donovan units three months post challenge. Giemsa stained smears of liver and spleen were microscopically observed to count the nuclei of parasite inside the macrophages. LDU was calculated as the number of amastigotes per 1000 cell nuclei multiplied by organ weight in grams (27).

### LDA

The viable parasite burden in the infected liver and spleen was estimated by LDA. Homogenized infected tissues were serially diluted, cultured, and monitored for the transformation of amastigotes into promastigotes. The LDA result was calculated based on the maximum dilution that led to parasite transformation (55).

### Statistical analysis

Prism-Graphpad version 5.0 software was used for analyzing the data. Data are represented as the mean ± standard error of mean. Two tailed Student’s *t*-test was used to analyze the difference within the groups and their significance. For group analysis, one-way ANOVA and post *t*-test analysis by Tukey’s multiple comparisons were employed. Statistical significance in the groups have been represented as * for *P*<0.05, ** for *P* <0.005 and *** for *P* <0.001.

## DECLARATIONS

### Ethics approval and consent to participate

All animal experiments were approved the institutional committee for animal ethics of CSIR- IICB under the guiding principle of CPCSEA, Government of India. Ethical Committee Approval number- IICB/AEC/Meeting/Feb/2018/10

### Consent for publication

Not applicable

### Availability of data and materials

All the information provided in the research is contained within the article. For additional information, please contact the corresponding author.

### FUNDING

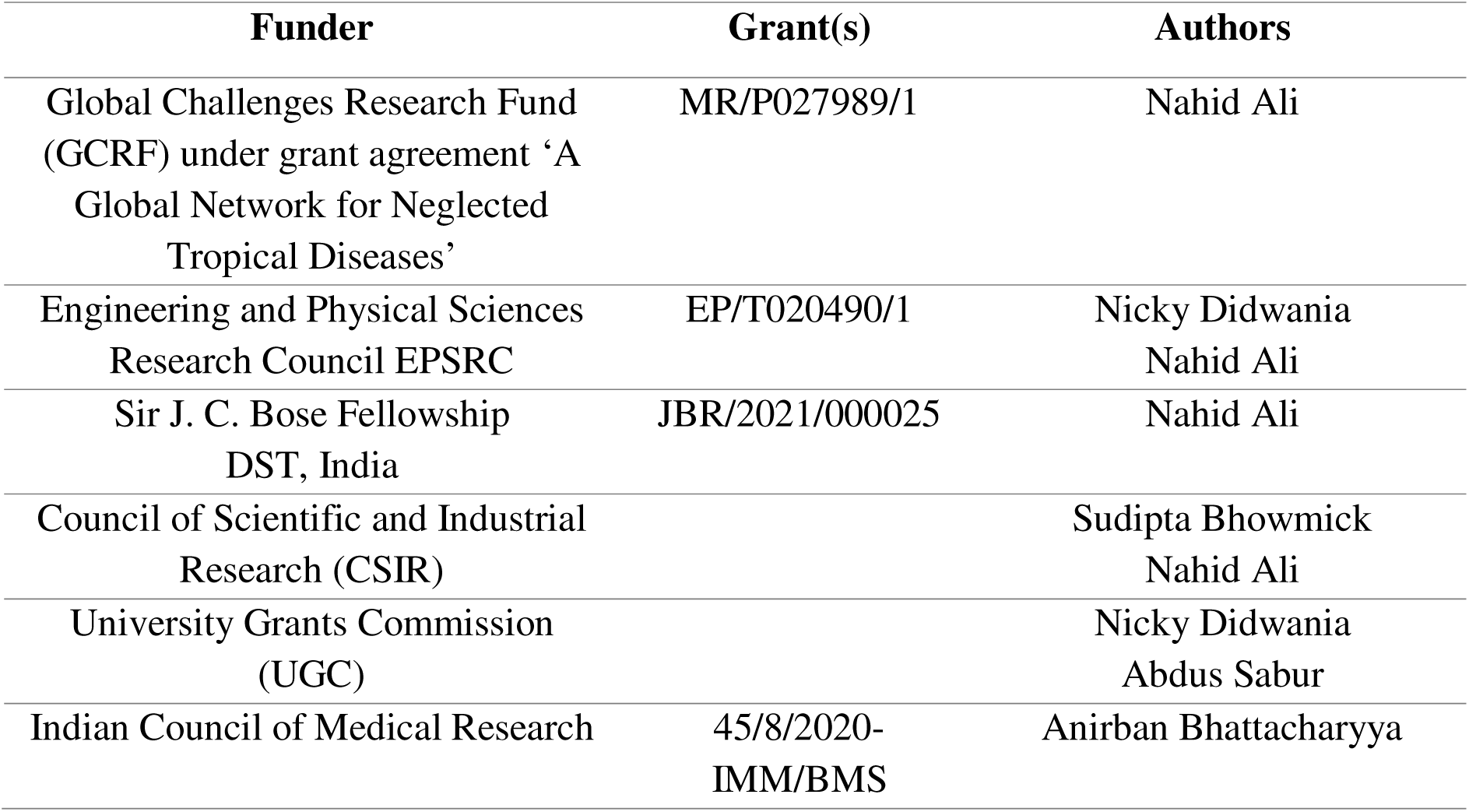

### Authors’ contributions

All the authors contributed significantly in conceptualising the study, designing experiments, sample collection, data acquisition, and analysis and interpretation of data. N.A. designed, analyzed, critically revised and approved the final version to be published. N.D. designed, performed and analyzed the experiments, prepared the figures and wrote the paper. S.B., A.S., and A.B. conceived, designed and performed the experiments. All authors have read and agreed to the published version of the manuscript.

## Acknowledgements

Not applicable

