## supplementary file for "Evaluation of *Leishmania* Homologue of Activated C Kinase (LACK) of *Leishmania donovani* in comparison to glycoprotein 63 as vaccine candidate against visceral leishmaniasis"

1. Infectious Diseases and Immunology Divison, Council of Scientific and Industrial Research (CSIR)-Indian Institute of Chemical Biology, 4, Raja. S.C. Mullick Road, Jadavpur, Kolkata -700028, India.
2. Present address: Dr. Kanailal Bhattacharyya College, 15, Kona Road, Ramrajatala, Santragachi, Howrah- 711104, India.
3. Present address: Riaganj Surendranath Mahavidyalaya Sudarshanpur, Raiganj, Uttar Dinajpur, Pin-733134, India.

Running Head: LACK vs. gp63: Evaluating Vaccine Candidates for VL

***To whom correspondence should be addressed:** Prof. Nahid Ali

**SUPPLEMENTARY MATERIALS**

**MATERIALS AND METHODS**

**Preparation of antigens**

Leishmanial membrane antigens (LAg) was prepared from stationary-phase promastigotes of *L. donovani* harvested after the third or fourth passage, following a previously described protocol. Briefly, stationary-phase promastigotes were washed three times with cold phosphate-buffered saline (PBS) and then resuspended in 5 mM Tris-HCl buffer (pH 7.6). The cell suspension was vortexed and subjected to centrifugation at 2,310 × g for 10 minutes to collect the membrane pellet. This crude ghost membrane pellet was further sonicated using an ultrasound probe sonicator (Misonix, Farmingdale, NY). The resulting suspension was centrifuged at 5,190 × g for 30 minutes, and the supernatant containing the leishmanial antigens was collected. The protein concentration was determined by the Lowry method, yielding approximately 5 mg of protein per gram of cell pellet. Sodium dodecyl sulfate polyacrylamide gel electrophoresis (SDS-PAGE) using a 10% gel was performed to verify the protein profile. Proteins were visualized with Coomassie brilliant blue staining.

**Electroelution of proteins from SDS-PAGE gels**

The LAg from *L. donovani* promastigotes was subjected to 10% SDS-PAGE and the proteins with molecular mass of 63-kDa and 34-kDa localized in gels, stained with Coomassie blue, were eluted by electrophoresis in running buffer (0.025 M Tris, 0.192 M glycine, 1% SDS) using an Electro-Eluter (model 422; Bio-Rad) at 10 mA for 5 h. After elution, the proteins were dialyzed, lyophilized and resuspended in PBS.

**Cloning of LACK gene**

Promastigotes of *L. donovani* strain AG83 (ATCC® PRA-413™) were cultured in M199 media containing 10% FBS and pelleted down. The genomic DNA of the parasites was isolated using genomic DNA isolation kit (Thermo Scientific) according to the company’s user manual. The genomic DNA from the parasites was used as DNA template to amplify the LACK gene from chromosome number 28. The sequence of LACK gene of *L. donovani* was obtained from the TriTrypDB genome database. Primers containing restriction sites for NcoI and HindIII were designed using Oligotech software. Codons for six histidine residues were added on the C terminus. The PCR reaction mixture containing genomic DNA, dNTPs, 10X dream Taq buffer, dream taq polymerase, forward primer, reverse primer, DMSO and nuclease free water was set up. The amplification of LACK gene was confirmed on 1% agarose gel electrophoresis. Primers used for amplification were as follows

Forward– 5’-AGAACCATGGCGCACCACCACCACCACCACAACTACGAGGGT CACCTGAAGGGCCACCGCGG -3’

Reverse – 5’- ATATAAGCTTTTAGTGGTGGTGGTGGTGGTGCTCGGCGTCGG AGATGGACCACACGCGG -3’

PCR conditions for amplification were as follows: one cycle of 10 mins at 95°C, 35 cycles of 30 sec at 95°C, 90 sec at 72°C, followed by final extension of 7 min at 72°C .

The PCR-amplified fragments were cloned into the bacterial expression vector pET28a (Novagen, Madison, WI, USA). Both pET28a and PCR-amplified fragments were double digested with Nco1 and HindIII restriction enzymes in the double digestion mixture. The double digested products were quantified by nanodrops (Thermo scientific, USA) and a ligation mixture of 20 µl was set which contained 140 µg of vector and 120 µg of the insert (1:4 molar ratio) with 1 µl of T4 DNA ligase and 2 µl 10 X ligase buffer. Ligation was carried out for 20 minutes at 22°C followed by heat inactivation at 65°C for 20 minutes. The ligation mixture was then incubated with competent DH5-α for 30 minutes followed by heat shock of 90 seconds at 42°C. The competent cells were allowed to revive for 1 hr in Super Optimal broth with Catabolite repression (SOC) media (Himedia, India). The cells were then plated in Luria Bertani (LB) Agar plate containing 50 µg/ml kanamycin. In order to determine the correct orientation of ligaton of the inserts and the vector, the colonies were subjected to colony PCR. The positive colonies were cultured in LB media and the plasmids were isolated from them. The plasmids were double digested with Nco1 and Hind III and checked on 1% agarose gel to determine the correct fallout. The plasmids which were found to have the fallout were selected for expression of LACK protein.

**Expression and purification of LACK proteins in Rosetta strain of *E. coli***

Plasmids isolated from positive clones were transformed in competent Rosetta strain of *E. coli* and plated on 40 µg/ml chloramphenicol and 50 µg/ml kanamycin containing LB agar plates. Log phase bacterial cultures were induced with IPTG (0.5 mM) and allowed to grow at 30̊ C for another 4 hrs before harvesting cells. The bacterial pellet obtained by centrifugation at 6000g for 10 mins were suspended in tris buffered saline-TBS (25 mM Tris-HCl, 300 mM NaCl), 1 mg/ml of lysozyme (Roche), and 1 mM phenylmethylsulfonyl fluoride (PMSF) pH 8.0. Bacterial cells were then sonicated and centrifuged to collect the inclusion bodies as pellet. Inclusion bodies were solubilised in binding buffer containing 8M Urea, 10 mM imidazole in 25 mM TBS and then centrifuged at 11,000 rpm for 30 mins. The supernatants were collected and added to pre-equilibrated Ni-NTA agarose matrix to allow His-tagged proteins to bind. The bound matrix was washed with wash buffer containing 8M Urea, 50 mM imidazole in 25 mM TBS with 0.1% Triton X 100. Finally the bound recombinants were eluted using elution buffer containing 8M Urea, 500 mM imidazole in 25 mM TBS. The purified proteins were dialyzed against TBS buffer with decreasing urea concentration and finally exchanged with 0.02M phosphate buffered saline (PBS), pH 7.2, in a 10-kDa Amicon cutoff membrane before analyzing the purity of LACK by SDS-PAGE.

**Entrapment of antigens in liposomes**

Cationic liposomes containing 63-kDa gp63, 34-kDa antigen and recombinant LACK were prepared with distearoyl derivative of L-α-phosphatidyl choline (DSPC), cholesterol (Sigma-Aldrich, USA) and stearylamine (Fluka, Switzerland) at a molecular ratio of 7:2:2. The lipid mixture was dissolved in chloroform at 60^0^C over a water bath. The solvent was then removed under reduced pressure by a rotary evaporator. The thin dry film was dispersed in 1ml PBS for the preparation of empty liposomes. Antigen entrapped liposomes were prepared by the dispersion of lipid film in 1ml PBS containing 1mg/ml of proteins. The mixture was vortexed and the suspension sonicated for 30 sec by an ultrasound probe sonicator (Misonix). Liposomes with entrapped antigens were separated from excess free antigen by three successive washing in PBS with ultracentrifugation (105,000 × g, 60 min, and 4^°^C). The amount of entrapped antigen was determined by the method of Lowry *et al.*, in the presence of 10% SDS.

**Cell proliferation**

The spleens were aseptically removed from the immunized, and 3 months infected BALB/c mice and single cell suspensions were prepared in RPMI 1640 supplemented with 10% FBS, l00 U/ml penicillin G sodium, 100 μg/ml streptomycin sulfate and 50 μM β-mercaptoethanol (Sigma-Aldrich, USA) (complete medium). RBCs were removed by lysis with 0.14 M Tris buffered NH_4_Cl. The remaining cells were washed twice with complete medium and viable mononuclear cell numbers were determined by counting Trypan blue unstained cells in a hemocytometer. Then the cells were cultured in triplicate in a 96 well flat bottom plate (Nunc, Naperville, IL) at a density of 2 ×10^5^ cells/well in a final volume of 200 μl and stimulated with either gp63 or 34-kDa antigen (2.5 μg/ml) in the presence or absence of anti-CD4 and anti-CD8 antibody (1 μg/10^6^ cells; BD Pharmingen). The cells were incubated for 96 h at 37^°^C in a humified chamber containing 5% CO_2_. Cells were pulsed with 1 μCi of [^3^H ]-Thymidine (Amersham Biosciences, Buckinghamshire, UK) per well 18 h before they were harvested on glass fiber paper. Thymidine uptake was measured in a β-scintillation counter (Beckman Instruments, Fullerton, CA). **Determination of antibody response by ELISA**

Serum samples from immunized and infected animals were analyzed by ELISA for the presence of antigen specific antibodies. In brief, 96-well microtiter plates (Maxisorp, Nunc) were coated with gp63 or 34-kDa antigen or LACK (10 μg/ml) diluted in PBS overnight at 4^°^C. The plates were blocked with 1% BSA in PBS at room temperature for 2 h to prevent nonspecific binding. After washing with PBS containing 0.05% Tween 20 (PBST) the plates were incubated overnight at 4^°^C with 1:2000 dilution of mice sera. Next day, the plates were again washed with PBST and incubated further for 1 h at room temperature with horseradish peroxidase conjugated goat antimouse IgG diluted at 1:3000 in blocking buffer. For isotype analysis, parallelly plates were incubated with horseradish peroxidase conjugated goat antimouse IgG1 or IgG2a (BD Pharmingen) at 1:3000 dilution for 1 h at room temperature. The plates were washed and 3,3',5,5'-tetramethylbenzidine (TMB) was added for 10 min and stopped with 2N sulphuric acid solution. The absorbance was read in an ELISA plate reader (Thermo, Waltham, MA) at 450 nm**.**

**Identification and estimation of cytokine producing lymphocytes by flow cytometry**

Splenocytes from immunized and infected mice were cultured in 24-well plates and stimulated with LACK in complete medium at 37°C. After 10-12 hrs of incubation at 37°C, Brefelden A was added to the cells, incubated further for 2 hrs at 37°C. Cells were then stained for CD3, CD4 and CD8. Following fixation and permeabilization, the cells were stained for cytokines, IFN-γ, IL-12, IL-2, IL-4, and IL-10. Fluorescence pattern of the stained cells were acquired by flow cytometry and analyzed by Flow Jo software as described previously.

**LDA**

To estimate live parasites in the infected liver and spleen, limiting dilution assay (LDA) was employed. For LDA, 1 mg homogenized infected tissues were five fold serially diluted in Schneider’s media contain 10% FBS and cultured for 3 weeks at 22°C days in 96-well culture plates allowing transformation to promastigotes. Each well was observed for the presence of promastigotes for 21 days. The log of five to the power of maximum dilution of tissue that led to transformation of amastigotes to promastigotes multiplied by the total organ weight in mg gave the estimate of viable parasite burden in the organs by LDA.
